# Genome-Epigenome Interactions Associated with Myalgic Encephalomyelitis/Chronic Fatigue Syndrome

**DOI:** 10.1101/237958

**Authors:** Santiago Herrera, Wilfred C. de Vega, David Ashbrook, Suzanne D. Vernon, Patrick O. McGowan

**Affiliations:** Centre for Environmental Epigenetics and Development, University of Toronto, Scarborough, ON, Canada; Department of Biological Sciences, University of Toronto, Scarborough, ON, Canada; Department of Cell and Systems Biology, University of Toronto, Toronto, ON, Canada; Solve ME/CFS Initiative, Los Angeles, CA, United States of America; Current affiliation: Department of Biological Sciences, Lehigh University, Bethlehem, PA, United States of America; Current affiliation: The Bateman Horne Center of Excellence, Salt Lake City, UT, United States of America

## Abstract

Myalgic Encephalomyelitis/Chronic Fatigue Syndrome (ME/CFS) is an example of a complex disease of unknown etiology. Multiple studies point to disruptions in immune functioning in ME/CFS patients as well as with specific genetic polymorphisms and alterations of the DNA methylome in lymphocytes. However, the association between DNA methylation and genetic background in relation to the ME/CFS is currently unknown. In this study we explored this association by characterizing the genomic (~4.3 million SNPs) and epigenomic (~480 thousand CpG loci) variability between populations of ME/CFS patients and healthy controls. We found significant associations of methylation states in T-lymphocytes at several CpG loci and regions with ME/CFS phenotype. These methylation anomalies are in close proximity to genes involved with immune function and cellular metabolism. Finally, we found significant correlations of genotypes with methylation phenotypes associated with ME/CFS. The findings from this study highlight the role of epigenetic and genetic interactions in complex diseases, and suggest several genetic and epigenetic elements potentially involved in the mechanisms of disease in ME/CFS.

## Introduction

Understanding the biological basis of complex traits and diseases remains one of the biggest challenges in biology and medicine. Chronic Fatigue Syndrome (also known as Myalgic Encephalomyelitis, hereafter referred to as ME/CFS) is an example of a complex, multifactorial disease with symptoms that vary substantially among patients. ME/CFS is a debilitating multisystem disease affecting between 1 and 2 million people in the United States alone^1^, with an annual economic impact between $17 and $24 billion ^2^. Yet its biological basis remains largely unknown.

Multiple studies point to disruptions in the immune and neuroendocrine systems in ME/CFS patients ^3–14^ ME/CFS appears to be associated with specific genetic polymorphisms ^15–17^, as well as with alterations of the DNA methylome in lymphocytes^14,18,19^. Understanding the contribution of the genetic background in ME/CFS patients as a predisposing factor for epigenetic abnormalities associated with the disease is a fundamental step to elucidate its causes. This understanding is also key for the development of tools to identify risk factors and potential treatments.

T-cell lymphocytes appear to be a primary cell type underlying immune and neuroendocrine abnormalities observed in ME/CFS patients. Functional impairment in T-cell glucocorticoid receptor and increased dexamethasone sensitivity are characteristic of some ME/CFS patients^14, 20^. Furthermore, genetic polymorphisms within non-coding regions of T-cell receptor loci^15^, as well as differential methylation in CD4+ T helper lymphocyte cells (Brenu et al., 2014), have been associated with the disease. The possible interactions between genomic and T-cell epigenomic variation in ME/CFS remain unknown.

In this study, we aimed to explore the association between DNA methylation profiles of T-cells and single nucleotide polymorphisms (SNPs) in ME/CFS patients. We quantified lymphocyte proportions and isolated CD3+ T-cells (including both CD4+ T helper cells and CD8+ T killer cells) via fluorescence activated cell sorting. We characterized the variation in genomic (~4.3 million SNPs) and epigenomic (~480 thousand CpG loci) variability among ME/CFS patients and healthy controls. Using this approach, we: 1) tested the association of genome-wide SNP genotypes with ME/CFS disease status; 2) tested the association of differentially methylated CpG loci and regions in CD3+ T-cells with ME/CFS disease status; 3) performed a methylation quantitative trait analysis to investigate the possible interactions between genetic background and methylation phenotypes of CD3+ T-cells associated with ME/CFS disease status.

## Methods

### Ethics approval and consent to participate

This study adhered to the human experimentation guidelines as outlined by the Helsinki Declaration of 1975. The collection of and analysis of clinical information and biological samples by the SolveCFS BioBank was ethically approved by the Genetic Alliance ethics review board (IRB # IORG0003358) and the University of Toronto (IRB #27391), which also approved all procedures for obtaining written informed consent from all participants in the study. Two copies of the consent form were signed, with one copy provided to the participants and one copy under secure storage at the SolveCFS Biobank.

### Study population

In total, 61 ME/CFS diagnosed patients receiving care at the Bateman Horne Center, Utah (46 females, 15 males) and 48 healthy control (36 females, 12 males) individuals were recruited for this study. Female to male ratios were nearly identical in both cases and controls (3:1). The sex ratio in our population of ME/CFS patients (cases) is consistent with previously reported ratios indicating predominance of this illness in females^21–23^. Diagnosis of ME/CFS was performed according to the 1994 Fukuda^24^ and 2003 Canadian^25^ criteria. To quantify functional impairment, individuals completed the standardized health-related quality of life survey RAND-36^26^. All individuals met the following criteria: 1) were HIV and Hepatitis-C negative; and 2) had no intake history of immunomodulatory or immunosuppressive medications. The Body Mass Index (BMI) of individuals ranged between 16.4 and 46 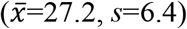, with no significant differences between case and control groups (t-test *p*=0.5; Wilcoxon *p*=0.74) (Fig. 1a). Similarly, age, which ranged between 18 and 62 years 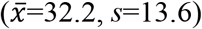, was not significantly different between cases and controls (t-test *p*=0.06; Wilcoxon *p*=0.07) (Fig. 1b).

**Figure 1.**
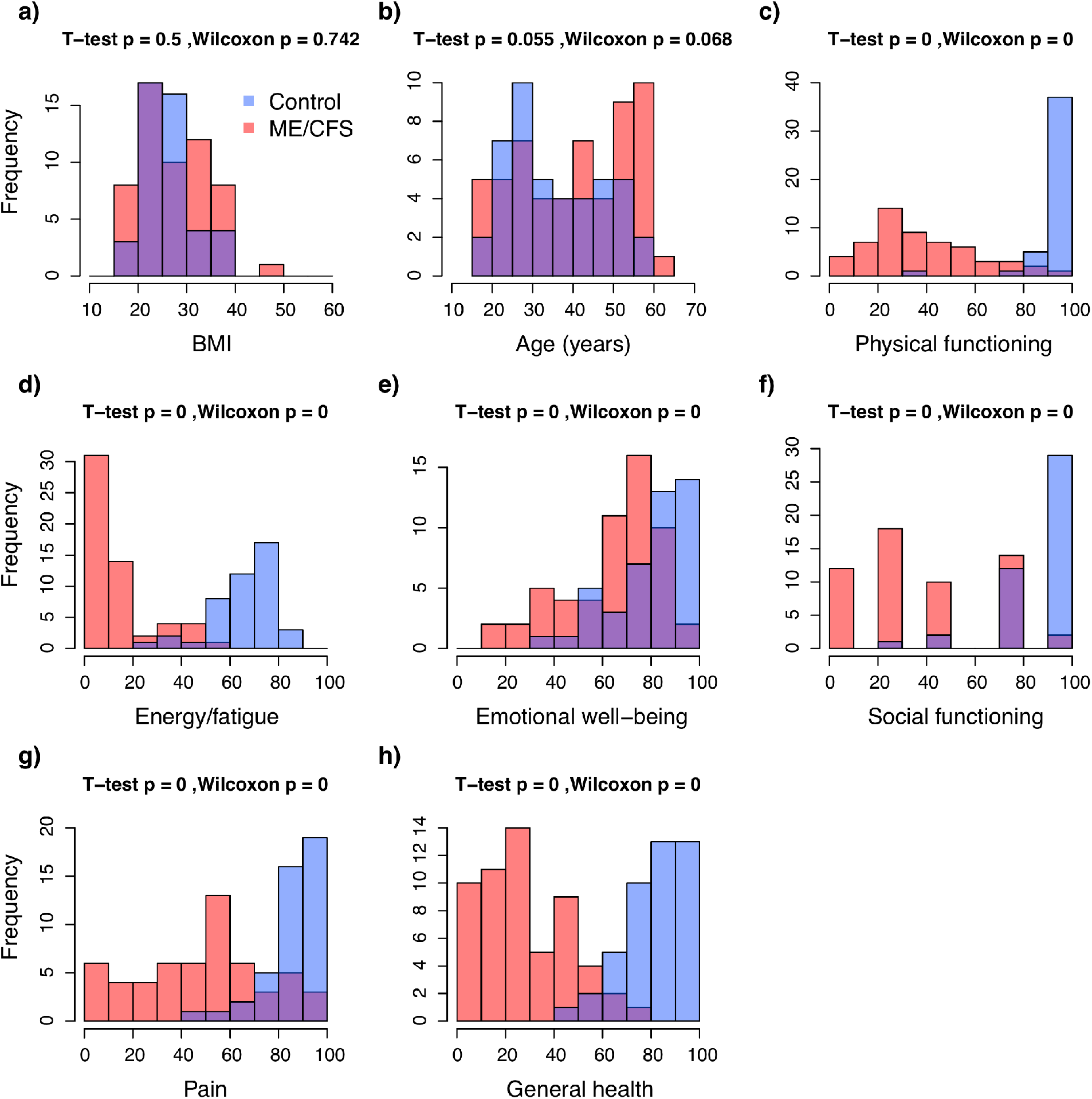
Frequency distributions of demographic and health-related indexes in the study population. **a**) Body mass index; **b**) Age; **c-h**) RAND-36 quality of life scales. Colours indicate the healthy control (blue, n = 48) and ME/CFS (red, n = 61) subpopulations. *p*-values from T-tests and Wilcoxon rank-sum tests.

### Sample processing

Whole blood from each individual was collected at the Bateman Horne Center and shipped overnight to Precision for Medicine, Maryland for Peripheral Blood Mononuclear Cells (PBMC) separation following procedures described in^14^ PBMCs were separated into two aliquots of approximately 7 million cells each, and shipped in liquid nitrogen to the Centre for Environmental Epigenetics and Development at the University of Toronto.

The first PBMC aliquot per patient was used for single nucleotide polymorphism (SNP) genotyping using the Human Omni 5-4 Array (Illumina Inc.). This array examines Single Nucleotide Polymorphisms at ~4.3 million loci throughout the human genome. Genomic DNA purification was performed with the MasterPure™ Complete DNA and RNA Purification Kit (Epicentre) following the standard protocol for cell samples. DNA quantity and purity was assessed using a NanoDrop 2000c Spectrophotometer (Thermo Scientific). Genotyping with the Omni 5-4 array was performed at the Princess Margaret Genomics Centre in Toronto.

The second PBMC aliquot was used for DNA methylation profiling of T-cells (CD3+) using the Human Methylation 450K Array (Illumina Inc.). This array quantifies methylation at ~480 thousand CpG loci throughout the human methylome. To quantify the relative proportions of cell type in the PBMC sample (i.e. CD4+ T-cells, CD8+ T-cells, CD19+ B-cells, and CD14+ monocytes) and isolate CD3+ T-cells, each sample was stained with fluorescently labelled antibodies and sorted in a FACSAria (BD Biosciences) flow cytometer at the Centre for Flow Cytometry and Microscopy of the Sunnybrook Research Institute in Toronto. Genomic DNA from T-cells was purified following the same procedure described above. Bisulfite conversion of purified DNA and CpG methylation profiling with the 450K Array was performed at the McGill University and Genome Quebec Innovation Centre in Montreal.

### Genome-wide association analyses

Analyses of SNP data quality and of association with ME/CFS disease phenotypes were performed with different parameters in the programs *GenABEL*^27^ and *PLINK*^28^, following standardized protocols^29, 30^. To minimize the number of false positive and negative associations, we first identified and excluded data from individuals that met one or more of the following criteria: 1) Inferred sex, as determined by the heterozygosity of the X chromosome, was incongruent with the known sex of the individual; 2) Heterozygosity rates or amount of missing data were outliers with respect to the whole population (Fig. 2a); 3) More than 10% of marker data was missing; 4) Relatedness to other samples, as measured by the identity by descent (IBD) statistic, was greater than that of a second- to third-degree relative (IBD>0.1875); 5) Ancestry, as determined by a principal components analysis (PCA) (Fig. 2b) performed with *EIGENSOFT*^31, 32^, was substantially different than that of the majority population in our cohort (i.e. European ancestry). Data from 10 individuals were excluded from all analyses: 4 cases (2 females, 2 males) and 6 controls (6 females). In addition to these criteria, we re-analysed the data excluding individuals with health-related quality of life measurements that overlapped between cases and controls. This exclusion of intermediate illness phenotypes was aimed at increasing the power to detect possible associations between disease status and (epi)genotypes by decreasing the heterogeneity in phenotype symptomatology within each group. We utilized the scores of RAND-36 scales (physical functioning, energy/fatigue, emotional well-being, social functioning, pain, and general health) as quantitative measurements of ME/CFS phenotypes because these were significantly different (α = 0.05) between case and control groups, prior to excluding individuals with intermediate phenotypes (Figs. 1c-h). RAND-36 scale scores were summarized into principal components (PC) using the *stats* package in R. We excluded data from case and control individuals with overlapping values along PC1 (Fig. 3), which explained ~80% of variance in the RAND-36 data. In total, data from 30 individuals were excluded using this approach: 18 cases (12 females, 6 males) and 12 controls (9 females, 3 males). These data were re-analysed in *PLINK.*

**Figure 2.**
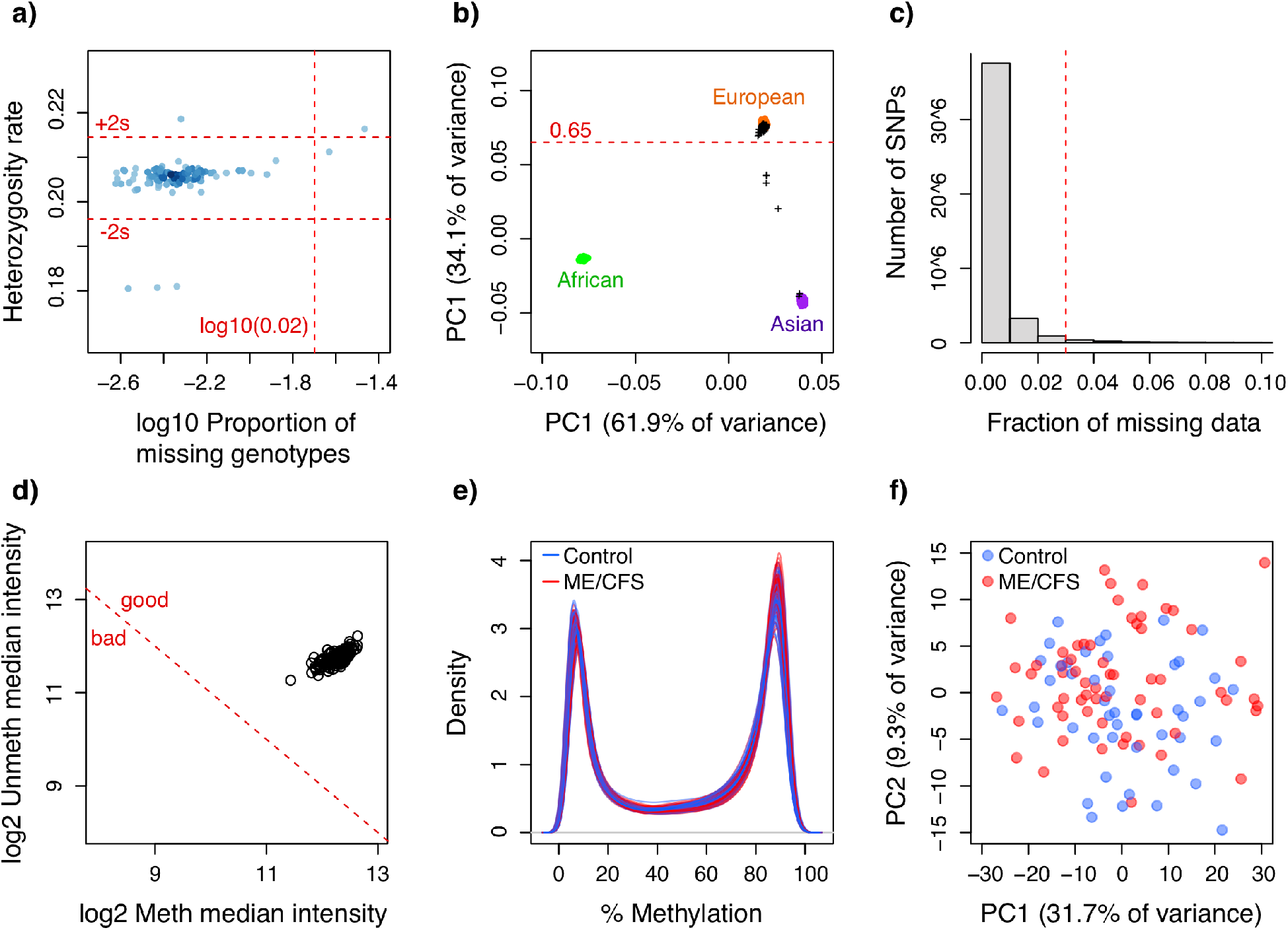
Quality control plots for SNP and CpG methylation data. **a**) Scatterplot of the proportion of missing genotypes vs. heterozygosity rate, per individual. Dot colour intensity indicates individual sample density. Horizontal red dotted lines indicate quality thresholds of ±2 standard deviations. Vertical red dotted line indicates a 2% missing data threshold; **b**) Inferred ancestry of individuals according to a principal components analysis of genotypes. The first two principal components are plotted. Genotyope data from individuals from reference populations (African, Asian and European) were obtained from the HapMap Phase III (HapMap3) database. Black crosses indicate individuals from this study. Horizontal red dotted line indicates European ancestry threshold; **c**) Frequency distribution of the fraction of missing data per SNP locus. Red dotted line indicates the 3% quality threshold; d) Scatterplot of median methylated and unmethylated fluorescence signals per individual. Dotted red line indicates quality threshold suggested in *minfi;* **e**) Methylation percentage (beta-values x 100) density distribution per individual; **f**) Scatterplot of the two principal components summarizing the variability in the methylation data per individual. For **e-f** colours indicate ME/CFS case (red, n = 61) or control (blue, n = 48) status of each individual.

**Figure 3.**
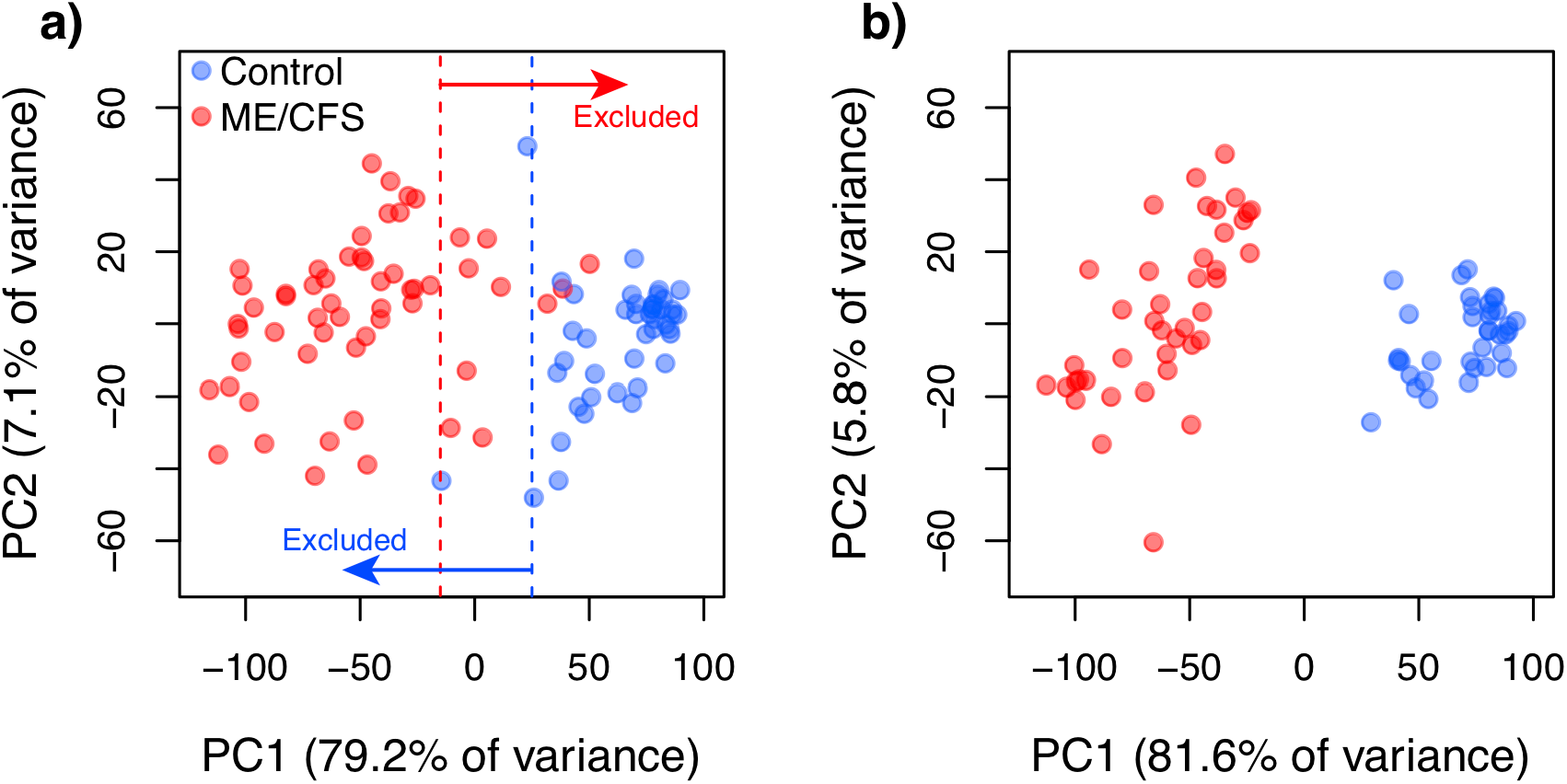
Scatterplots of the two principal components summarizing the variability in the standardized health-related quality of life surveys (RAND-36) per individual. **a**) Before exclusion of data from individuals with intermediate (overlapping) phenotypes along PC1. Dotted lines indicate the thresholds used to define intermediate phenotypes; **b**) After exclusion of data from individuals with intermediate phenotypes. Colours indicate the healthy control (blue, **a**) n = 48; **b**) n = 36) and ME/CFS (red, **a**) n = 61; **b**) n = 43) subpopulations.

After excluding individuals with sub-optimal data, we identified and excluded data from SNP loci that met one or more of the following criteria following^29, 30^: 1) Rate of missing genotypes was greater than 3% in *PLINK* (5% in *GenABEL)* (Fig. 2c); 2) Rate of missing data was significantly different *(p* < 0.00001) between cases and controls; 3) Allelic frequencies significantly deviated from Hardy-Weinberg equilibrium (χ^2^ *p* < 0.00001 in *PLINK,* FDR < 0.2 in *GenABEL);* and 4) Minimum allele frequency was smaller than 1% in *PLINK* (2% in GenABEL). Out of 4,284,426 genotyped SNP loci, 1,779,031 SNP loci were excluded from *PLINK* analyses, and 2,142,548 from *GenABEL* analyses.

We examined possible associations between SNP genotypes and ME/CFS disease phenotypes (case or control) using χ^2^ testing as well as logistic regression tests that included covariates of age, sex and BMI in both *PLINK* and *GenABEL.* To account for the uncertainty in the potential genetic model of inheritance of ME/CFS, we performed multiple tests with different underlying models: Genotypic, dominant, recessive, Cochram-Armitage trend, and allelic for the simple χ^2^ tests; and genotypic, dominant, recessive and multiplicative for logistic regressions. No inflation of test statistics was observed in any test (λ ranged between 1 and 1.01). To assess the significance of associations we: 1) Adjusted raw p-values for multiple testing following the Bonferroni^33^, Holm^34^, and Benjamini and Hochberg false discovery rate (FDR) corrections^35^; and 2) Calculated corrected (empirical) p-values (family wise) after 10,000 permutations. We generated spatial visualizations of raw p-values for all associations across chromosomes using the program *Haploview*^36^. Due to the higher prevalence of ME/CFS in females than males^21–23^, association tests were also performed in data from females only.

### SNP characterisation

SNPs with significant associations were examined using the following reference tools: the NHGRI-EBI catalogue of genome-wide association studies (<u>http://www.ebi.ac.uk/gwas/home</u>;^37^, the Ensembl genome browser (<u>http://www.ensembl.org/Homosapiens/Info/Index</u>;^38^, the Single Nucleotide Polymorphisms Annotator SNiPA (<u>http://snipa.helmholtz-muenchen.de/snipa3/</u>;^39^, the Genotype-Tissue-Expression database GTEx (<u>https://www.gtexportal.org/home/</u>;^40^, the genome-wide association study of blood plasma proteome database pGWAS (<u>http://metabolomics.helmholtz-muenchen.de/pgwas/</u>;^41^, and the developing brain methylation quantitative trait loci database (<u>http://epigenetics.essex.ac.uk/mQTL/</u>;^42^.

### Re-analysis of published GWAS data

Data from the ME/CFS genome-wide association study (GWAS) by Schlauch et al. (2016), were acquired from dbGAP (phs001015.v1.p1), and re-analysed following the pipeline described above to identify commonalities. The *crlmm* R Bioconductor package^43, 44^ was used for genotyping, and analysed in *GenABEL,* using the same thresholds as above. 704,464 SNP markers from 66 subjects passed our quality controls (44 females and 22 males, 36 cases and 30 controls). Because of the different arrays used (Illumina Human Omni 5-4 Array in this study vs. Affymetrix Genome-Wide Human SNP Array 6.0 used by Schlauch et al.) we constructed linkage disequilibrium (LD) proxies using *LDlink*^45^, with R^2^ ≥0.8, to make the results comparable.

### Gene-set analysis

The program *MAGMA*^46^ was used to complete a generalized gene-set analysis of the SNP data. This analyses focuses on genetic associations with phenotype at the level of genes and gene-sets rather than individual SNPs. This strategy augments the power to detect associations with complex traits and diseases, such as ME/CFS. Gene sets were taken from Molecular Signatures Database (MSigDB)^47^, including hallmark gene sets (hallmark gene sets summarize and represent specific well-defined biological states or processes and display coherent expression;^48^), canonical pathways (gene-sets from KEGG, BioCarta and Reactome) and GO gene sets (gene-sets that contain genes annotated by the same gene ontology term).

### Epigenome-wide association analyses

Analyses of CpG methylation data quality and of association with ME/CFS disease phenotypes were performed in the R package *minfi*^49^, following the pipeline suggested by^50^. All individuals identified for exclusion during the genome-wide association analyses were also excluded from this dataset to increase the power of detection of possible associations. In addition, we attempted to identify data from individuals that could represent potential outliers by the following graphical approaches (as suggested by^49^): 1) Comparing inferred sex versus known sex; 2) Examining intensity distributions of control CpG probes; 2) Plotting median methylated and unmethylated fluorescence signals; 3) Plotting methylation percentage density distributions; and 4) Summarizing the variability in the methylation data through a principal component analysis. No individuals were identified as outliers (Figs. 2d-f). We discarded data from CpG loci that: 1) Contained known SNPs at the methylation dinucleotide; or 2) Contained missing data. Raw probe florescence intensities were normalized by Subset-Quantile Within Array Normalization^51^, which takes into account the differences between Infinium type I and II probes. The level of methylation in each CpG locus was measured as beta-values, ranging from 0 to 1, which represent the proportion of methylation. In total, 467,971 CpG loci (out of 485,512) were retained for further analyses.

We examined possible associations between methylation levels at CpG loci and ME/CFS disease phenotypes (case or control) through F-tests of logit-transformed beta-values^52^ as implemented in the *dmpFinder* function. To correct for potential confounding effects of multiple methylation array batches (two in this study), as well as covariates of age, sex and BMI, we utilized the empirical Bayes procedure implemented in the R package *ComBat*^53^. To assess significance of associations (α = 0.05) we: 1) adjusted raw p-values for multiple testing by performing Benjamini and Hochberg false discovery rate (FDR) corrections; and 2) calculated empirical p-values after 10,000 permutations as described in^14^. In addition to testing for associations at individual CpG loci, we performed tests of association at differentially methylated genomic regions using the R package *bumphunter* as described in^54^. Genomics regions were defined as clusters of CpG loci within a 500bp region. We assessed the significance of associations (α = 0.05) by calculating empirical p-values from null distributions of test statistics after 1,000 bootstrap pseudoreplicates.

### Genome-epigenome association analyses

To identify associations between SNP genotypes and methylation levels at significantly differentially methylated CpG loci, we performed a methylation quantitative trait loci (mQTLs) analysis using linear additive regression models (including covariates) in the R package *Matrix eQTL*^55^. Only local cis-mQTL were considered, i.e. CpG-SNP pairs that are within 1Mbp of each other. Both CpG loci and SNPs were mapped to the UCSC human genome assembly version hg19 (Genome Reference Consortium GRCh37)^56^. mQTLs were considered significant when FDR corrected p-values were smaller than α = 0.05.

### Enrichment analysis

We carried out gene-set enrichment analyses for genes of interest. The R package *clusterProfiler* 3.4.4^57^ was used for Gene Ontology Biological Process (GO BP)^58, 59^ and KEGG pathway^60, 61^ enrichment analysis, with *p* and *q* value cut-offs of ≤ 0.05. Reactome pathway analyses were carried out using the *ReactomePA* 1.20.2 R package^62^, with a p-value cut-off of ≤ 0.05. All three packages reference the latest versions of their respective databases. These significantly enriched annotations were visualized with the ‘enrichMap’ function of *DOSE* 3.2.0 R package^63^, with specific parameters to aid legibility using different numbers of enriched annotations.

## Results & Discussion

### Lymphocyte proportions

In contrast to the significant differentiation in health-related quality of life RAND-36 scales between ME/CFS cases and healthy controls (Fig. 1c-h), there were no differences in the relative proportions of cell types in the PBMC lymphocyte samples (Fig. 4). Previous studies investigating potential differences in the relative proportions of general lymphocyte types in ME/CFS patients have produced incongruent results which have ranged from an increased proportion of CD8+ T cells (Klimas et al., 1990) to no significant differences^7, 64^ These observations suggest that alterations of general lymphocyte type proportions may not be a characteristic feature of ME/CFS. Rather, abnormalities in immune system functioning associated with ME/CFS appear to involve alterations in the activity and abundance of specific sub-populations^5,6,65^.

**Figure 4.**
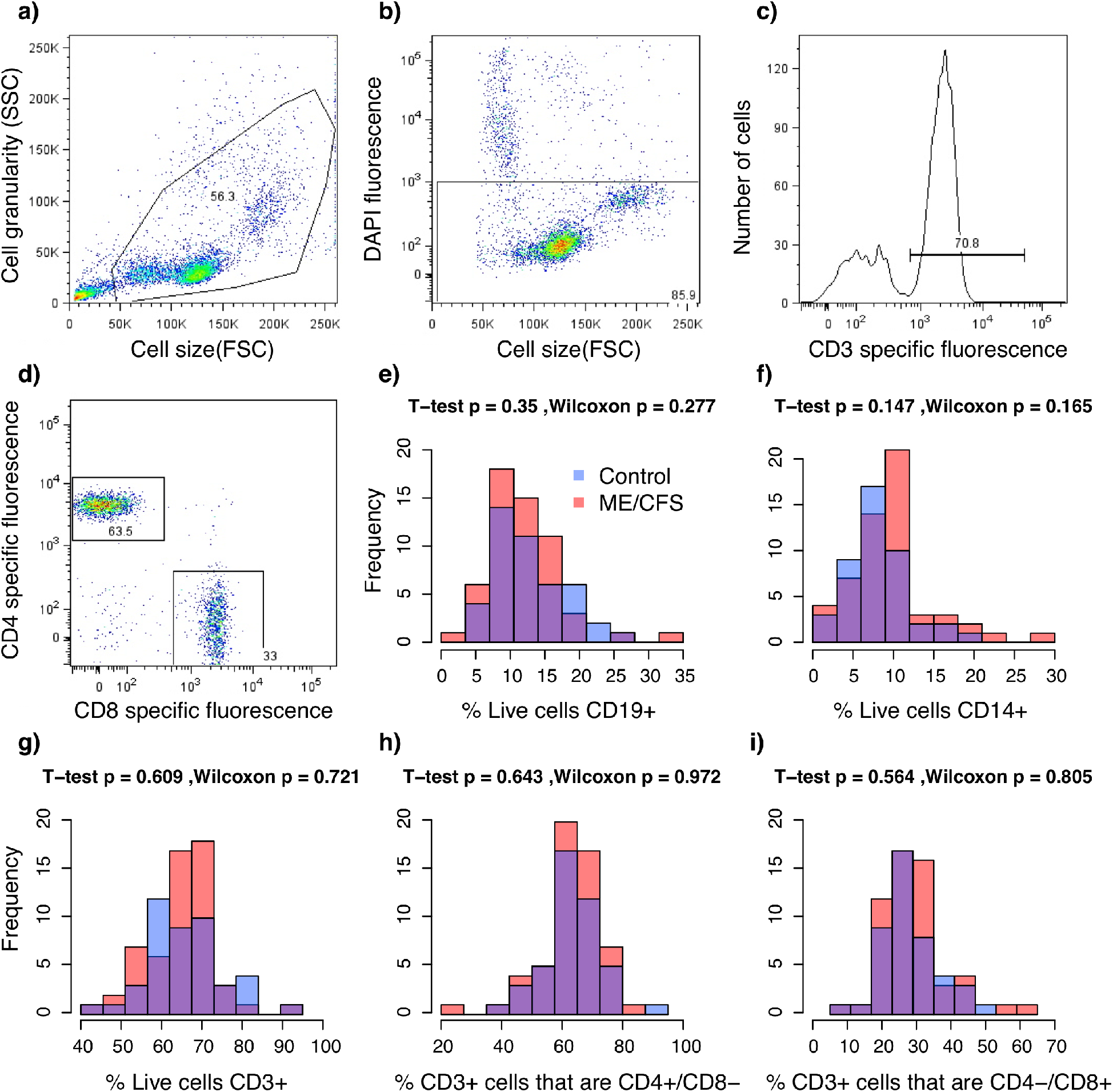
Results from florescence-activated cell sorting (FACS) of PBMCs. **a-d** Representative example of sorting parameters from one individual. **a**) Total particle composition of sample before gating; **b**) Gated cells showing live cells in rectangle; **c**) Frequency of gated T-cells (CD3+); **d**) CD4/CD8 expression on CD3+ gated cells. **d-i** Frequency distributions of relative proportions of cell types per individual. Colors indicate the healthy control (blue, n = 48) and ME/CFS (red, n = 61) subpopulations. *p*-values from T-tests and Wilcoxon rank-sum tests. **e**) CD19+ B-cells; **f**) CD14+ monocytes; **g**) CD3+ T-cells; **h**) CD4+/CD8-T-cells; **i**) CD4-/CD8+ T-cells.

### Genetic associations with ME/CFS

None of the more than 2 million variable SNP loci targeted in this study had a significant association (α = 0.05) with ME/CFS after p-value corrections with Bonferroni, Holm, Benjamini and Hochberg, or permutation methods when data from both sexes were analysed together. This result was consistent across all the χ^2^ and logistic regression tests (Figs. 5a-b summarize the results of the simple χ^2^ genotypic test as a representative example). Because of the known increased prevalence of ME/CFS in females^21–23^, we performed independent analyses of data from females only. These analyses revealed a significant association (χ^2^ genotypic test, permutation-corrected p-value = 0.0374, OR = 0.1845, 1/OR = 5.42) of one SNP (rs11712777, chr3:42347678) with the ME/CFS disease phenotype.

**Figure 5.**
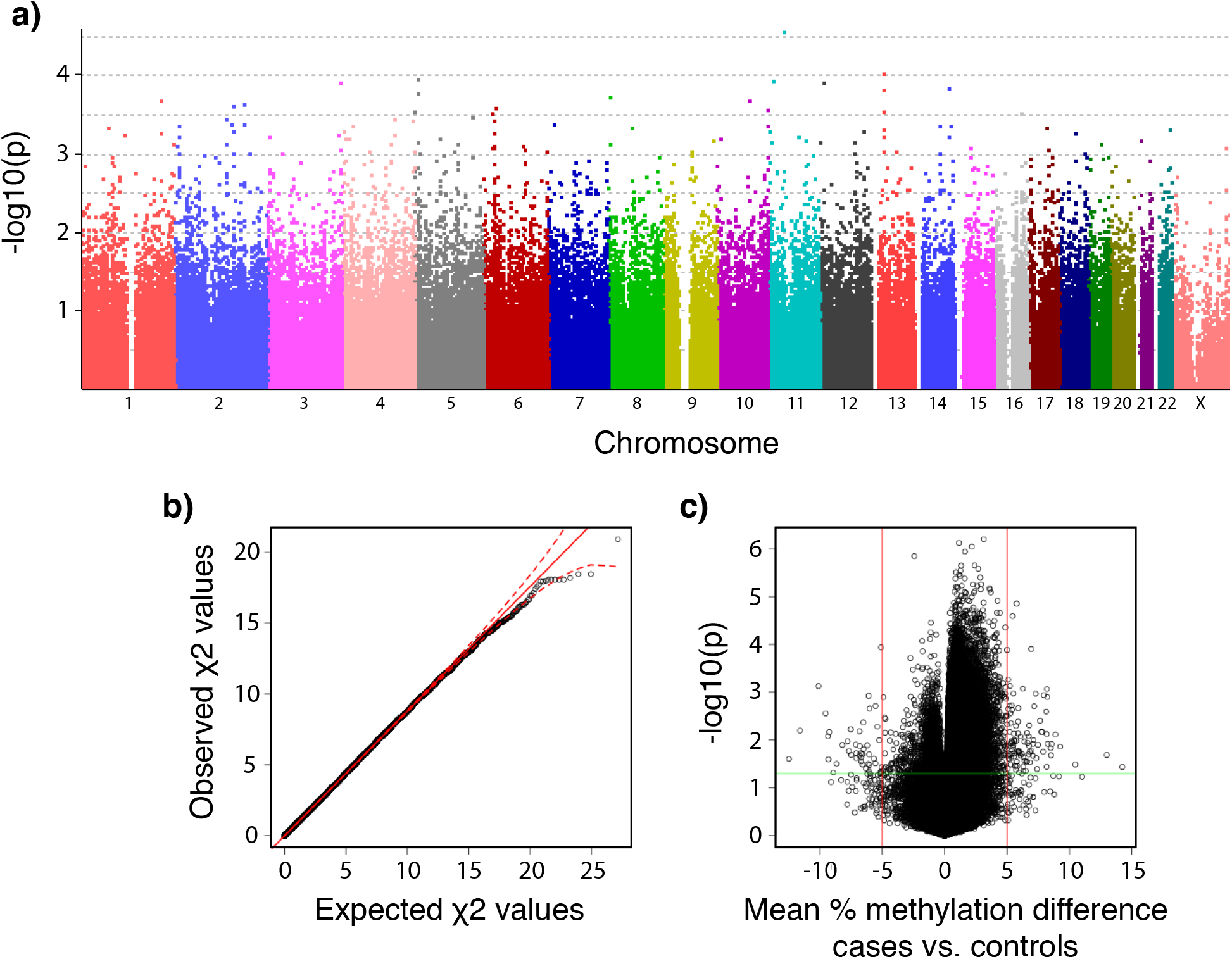
Plots summarizing the strength of associations between SNP genotypes and DNA methylation levels to disease phenotypes (healthy controls, n = 36; vs. ME/CFS cases, n = 43) in data from males and females subpopulations. **a**) Manhattan plot of p-values calculated from the simple χ^2^ genotypic test of association for 2,505,395 SNPs *(PLINK* analysis). Currently accepted genome-wide significance threshold is 5x10^−8^ (7.3 in –log10 units). Bonferroni’s adjustment significance threshold for this study is 2x10^−8^ (7.7 in -log10); **b**) Quantile-quantile plots of expected vs. observed χ^2^ test statistics from the simple χ^2^ genotypic test of association. Red solid line indicates the middle of the first and third quartile of the expected distribution of the χ^2^ test statistics. Red dashed lines indicate the 95% confidence intervals of the expected distribution of the χ^2^ test statistics; c) Volcano plot of effect size (mean percentage DNA methylation difference between ME/CFS and controls) vs. association empirical p-values calculated after 10,000 random permutations. Vertical red lines indicate biological significance threshold of 5% absolute difference in methylation at each locus. Horizontal green line indicates statistical significance threshold of p <0.05.

These results are contrasting to previous genotype association analyses in ME/CFS populations, which have found statistically significant associations in multiple loci. The earliest study by Smith et al. (2011) evaluated 116,204 SNPs (n=40 CFS, n=40 non-ME/CFS) using the Affymetrix GeneChip Mapping 100K array, and found 65 SNPs associated with ME/CFS (p<0.001). Rajeevan (2015) used the Affymetrix Immune and Inflammation Chip to focus on ~11,000 SNPs located in genes involved in immune and inflammation pathways (n=121 ME/CFS, n=50 non-ME/CFS). Of these, 32 were associated with ME/CFS (p<0.05). Most recently, Schlauch et al. (2016) evaluated 906,600 SNPs with the Affymetrix Genome-Wide SNP Array 6.0 (n=42 ME/CFS, n=38 non-ME/CFS) and found 442 SNPs that were associated with ME/CFS (P<3.3×10-5). The SNP that we found in significant association with ME/CFS in females, rs11712777, was not included in any of these datasets. One SNP in the Schlauch et al. (2016) data, rs1468604, is in linkage disequilibrium (LD) with rs11712777 (r^2^ = 0.8716; European population). The apparent discrepancy could be explained by the imperfect linkage between the two SNPs, and therefore we recommend rs11712777 as a candidate for direct genotyping in future studies.

There are no other overlaps in the SNPs or genes associated with ME/CFS between this study and previous genetic association studies. This observation may be confounded by a combination of multiple factors, including: 1) Differences in the types of arrays utilized in each study (our study, with the largest genetic coverage to date, evaluated two-orders of magnitude more SNPs than the Rajeevan (2015) study); 2) Differences among cohorts due to the wide heterogeneity of ME/CFS; 3) Reduced statistical power to discriminate the effects of multiple small-effect variants due to relatively small sample sizes; and 4) Interactions with environmental and epigenetic factors. Additional larger-scale genome-wide association studies with overlapping SNP probes and larger sample sizes will further our understanding of the interaction between genetic factors and ME/CFS.

Generalized gene-set analysis of the SNP data in *MAGMA* did not identify any gene-set significantly enriched in either our data, or the data from Schlauch et al. (2016).

### Characterisation of SNP rs11712777

We used a variety of online reference resources to characterize the current knowledge of rs11712777, and how it may influence ME/CFS phenotype (see Methods section). We also examined SNPs in high LD with SNP rs11712777 (R^2^ ≥0.8; Table 1). The Genotype-Tissue-Expression database (GTEx) indicates that rs11712777, and the genes in LD with it, form an expression quantitative trail loci (eQTL) altering the expression of the CCK (cholecystokinin peptide hormone) gene. CCK has a number of active forms, expressed in a variety of tissues, including the blood, intestine and blood^66^, and plays a role in appetite, body weight and the immune system^66, 67^ A rat knockout (KO) of the cholecystokinin B receptor (CCKBR) shows attenuated sickness behaviour^68^. This sickness behaviour in rats has remarkable similarity to some of the symptoms of ME/CFS^69^, including fatigue, malaise, hyperalgesia, sleepiness, anhedonia, weight loss and diminished activity^69^. CCK is also co-localized with sleep-promoting preoptic neurons in the hypothalamus^70^, which may relate to fatigue and unrefreshing sleep symptoms in ME/CFS. Finally, recent evidence suggests that CCK has a role regulating the differentiation of CD4+ T-cells^71^, and that CCK-expressing neurons are a critical cellular component of the hypothalamic-pituitary-adrenal axis^72^. These roles of CCK in components of the immune system are consistent with suggested immune dysregulation in ME/CFS^3–8,14^ While CCK-associated variant rs11712777 may be a biologically relevant candidate influencing susceptibility to ME/CFS, our findings suggest that it only accounts for a small fraction of the risk (OR = 0.1845). However, it constitutes a relevant target for future research.

**Table 1.** SNPs in high LD (R2 ≥0.8) with candidate SNP rs11712777. MAF = Minor allele frequency. Adapted from <u>https://analysistools.nci.nih.gov/LDlink/</u>.

| RS number | Coordinate | Alleles | MAF | Distance | D' | R<sup>2</sup> | Correlated alleles |
| --- | --- | --- | --- | --- | --- | --- | --- |
| rs11712777 | chr3:42347678 | (C/T) | 0.3877 | 0 | 1 | 1 | C=C,T=T |
| rs17223780 | chr3:42363369 | (C/T) | 0.3598 | 15691 | 0.9955 | 0.8799 | C=C,T=T |
| rs11715412 | chr3:42368008 | (A/G) | 0.3608 | 20330 | 0.991 | 0.8757 | C=A,T=G |
| rs1966393 | chr3:42368673 | (G/A) | 0.3608 | 20995 | 0.991 | 0.8757 | C=G,T=A |
| rs17224501 | chr3:42369441 | (G/A) | 0.3608 | 21763 | 0.991 | 0.8757 | C=G,T=A |
| rs1468604 | chr3:42368882 | (T/C) | 0.3618 | 21204 | 0.9865 | 0.8716 | C=T,T=C |
| rs35392307 | chr3:42372207 | (G/A) | 0.3579 | 24529 | 0.9909 | 0.8643 | C=G,T=A |

In addition to rs11712777, SNP rs17223780 (R^2^ = 0.8799) binds DNase in CD14+ monocytes (<u>http://www.regulomedb.org/snp/chr3/42363368</u>), indicating a possible regulatory role in the immune system. Another SNP in the vicinity of rs11712777 (D’ = 0.7211, R^2^ = 0. 0126), rs33449 (chr3:42400801), is associated with increased daytime resting duration (<u>http://www.ebi.ac.uk/gwas/search?query=3:42347678-42372207</u>;^73^). This is a phenotype that may be related to the fatigue aspect of ME/CFS.

### Epigenetic associations with ME/CFS

Of the 467,971 CpG loci analysed, 141 had significant associations with the ME/CFS phenotype (raw p-value < 0.05) and a mean percentage methylation difference between cases and controls greater than 5% when data from both sexes were analysed together (Fig. 5c). None of these differentially methylated loci were significant after FDR corrections, however 133 had significant empirical p-values < 0.05 calculated through permutation analyses (these are referred to as differentially methylated probes -DMPs; Supplementary Table S1). Analyses of methylation data from females alone indicated that 108 CpG loci had significant associations with the ME/CFS phenotype (raw p-value ≤ 0.05) and a mean percentage methylation difference between cases and controls greater than 5%. None of these differentially methylated loci were significant after FDR corrections, however 94 DMPs had significant empirical p-values ≤ 0.05 after permutation analyses (Supplementary Table S2). Out of these 94 DMPs, 29 were common to the DMPs found when analysing the data from both sexes combined.

Approximately half of the DMP were clustered in differentially methylated regions (DMRs). We found 17 DMRs with significant association with the ME/CFS phenotype (p-value < 0.05) when data from both sexes were analysed together (Supplementary Table S3). There were 22 DMRs when only females were considered (Supplementary Table S4). All of these regions were located nearby genes. 5 DMRs were found upstream of genes, 3 in promoters, 3 overlapping the 5’ end and 1 the 3’ end of genes, 10 inside introns, and 3 downstream of genes. DMR length, in terms of number of CpG loci ranged between 2 and 10 (average 2.72). 7 DMRs containing CpG loci identified as DMPs were detected in common between analyses of methylation data from both sexes and from females only (Figure 6).

**Figure 6.**
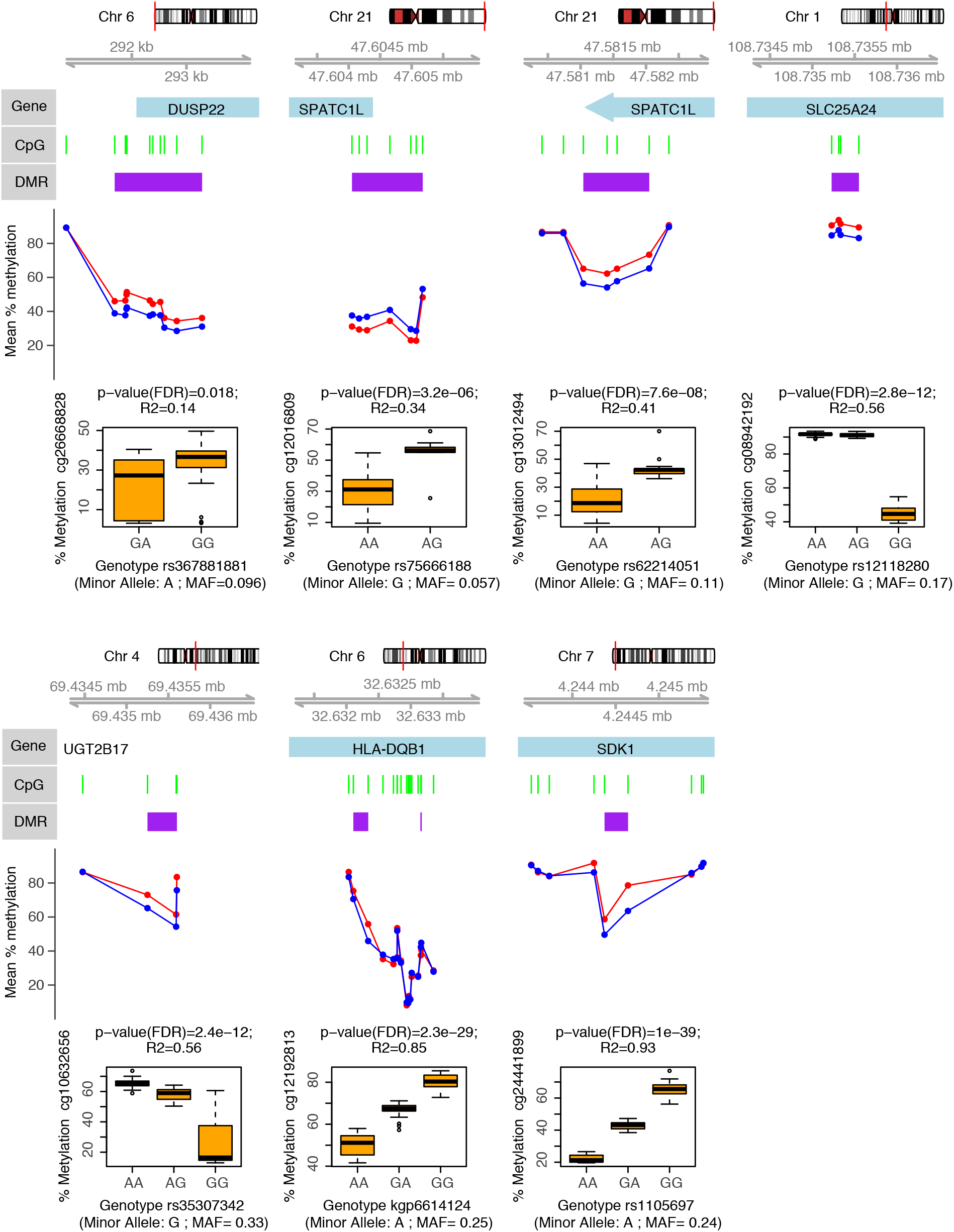
Genes associated with differentially methylated regions (DMR) in ME/CFS. Figure shows DMP-containing DMRs identified from the analysis of methylation data from both sexes (healthy controls, n = 36; vs. ME/CFS cases, n = 43) that were also identified from analysis of data females alone (healthy controls, n = 27; vs. ME/CFS cases, n = 34). Each panel shows (in descending order): 1) The chromosomal location of the gene/DMR; 2) The position of the DMPs (green bars) and DMR (purple bars) with respect to the gene (blue bars); 3) The mean percentage methylation difference between ME/CFS cases (red) and controls (blue) at each DMP; 4) The most significant meQTL association (as indicated by the R^2^ and p-values) between SNP genotype and the individual percentage methylation at the most significant DMP (as indicated by the p-value) within each DMR.

These results are in contrast with previous findings by our group, which revealed thousands to tens of thousands of differentially methylated CpG loci associated with ME/CFS in PBMCs, using the same 450K array^14, 18^. It is possible that differences between the targeted cell populations (i.e. PBMCs vs. isolated T-cells) may have contributed to the differences in the number of differentially methylated CpGs. The number of cell types within PBMCs may broaden the spectrum of epigenetic marks and thus increase the number of possible associations with the ME/CFS disease phenotype. Consistent with this idea, Brenu et al (2014) found 120 differentially methylated CpGs associated with ME/CFS in CD4+ T-cells (p<0.001) using the 450K array (n=25 ME/CFS, n=18 non-ME/CFS). This number of differentially methylated CpGs is similar to the 133 DMPs we found in this study, which targeted a broader T-cell population (including CD4+ and CD4-T-cells). However, the only overlap between our study and the study from Brenu et al (2014) corresponded to the HLA-DQB1 (major histocompatibility complex, class II, DQ beta 1) gene. HLA-DQB1 encodes a protein that is part of the DQ heterodimer, a cell surface receptor that is essential in immune signalling. We found two contiguous differentially methylated regions within an intron of this gene (Supplementary Tables S1 and S3, Figure 6). One region was hypermethylated whereas the other was hypomethylated in the ME/CFS group. Interestingly, the gene HLA-DQB1 contained cis-mQTLs significantly associated with these two DMRs (see next section, as well as Supplementary Table S5, and Figure 6). Brenu et al. (2014) found CpG hypermethylation associated with the HLA-DQB1 gene, however the specific location of this association was not reported. Recent studies focusing on CD4+ T-cells of patients affected by immune disorders such as rheumatoid arthritis^74^ and multiple sclerosis^75^ have found differential methylation in HLA-DQB1. This result is consistent with a potential immune dysregulation in ME/CFS.

We found 31 genes associated with DMPs in T-cells that were common to this study and a previous study by our group^14^ These genes, which include PAX8 (paired box 8), and ATP4B (ATPase H+/K+ transporting beta subunit) (Supplementary Table S1), are involved in the regulation of cellular processes and cell signaling. This is in line with recent ME/CFS work that observed differences in cellular metabolism in ME/CFS^76–78^.

Our results suggest that DNA methylation modifications in T-cells in ME/CFS are associated with the cellular metabolism differences that are observed in the disease and may play a role in the development of these phenotypic differences, however future work is required to understand this relationship.

### Genetic and epigenetic interactions associated with ME/CFS

All the DMPs identified according to empirical *p*-values had significant associations (FDR corrected p-values < 0.05) with SNP genotypes (independent of disease phenotype). In total there were 13,060 significant cis-mQTLs (Supplementary Table S5). Figure 6 shows the strongest SNP-DMPs cis-mQTLs associations (according to correlation coefficient R^2^) in each of the 7 DMP-containing DMRs that were common in analyses of methylation data from both sexes and from females only.

SPATC1L (spermatogenesis and centriole associated 1 like) and DUSP22 (dual specificity phosphatase 22) were the two genes containing cis-mQTLs with the largest differentially methylated regions: 11 DMPs (7 hypermethylated probes in 5’ UTR and 4 hypomethylated probes in 3’ UTR region) in SPATC1L and 10 hypermethylated probes in the 5’ UTR of DUSP22 (Supplementary Table S3 and Figure 6). While the exact function of SPATC1L is not well understood, it has been previously associated with xenobiotic response and differential methylation in the promoter of this gene is characteristic of certain ethnic groups in human populations^79^. DUSP22 hypermethylation has also been observed in the 5’ UTR region in T-cells of rheumatoid arthritis patients^80^. In T-cells, DUSP22 is known to inhibit proliferation and autoimmunity through inactivating Lck and preventing the activation of the T-cell receptor^81^. However, it remains to be confirmed how hypermethylation in the 5’ UTR region affects the overall activity of DUSP22 in T-cells of ME/CFS patients.

These results suggest that ME/CFS patients have differential methylation patterns in T-cells that are strongly correlated with the underlying genotype. Understanding the mechanisms of these interactions is a promising direction of research in ME/CFS.

## Conclusions

We identified over one hundred differentially methylated CpG loci associated with ME/CFS in T lymphocytes. Approximately half of these were clustered in differentially methylated regions of 500bp in size or less. Our data and analyses suggest that there is an indirect role of genotype influencing DNA methylation patterns associated with ME/CFS. We found no substantial large-effect direct associations of specific genotypes with ME/CFS disease phenotype. Larger scale genome wide association studies are necessary to test for potential small-effect associations between genotype and ME/CFS phenotype.

All of the methylation values at differentially methylated loci in T lymphocytes had significant correlations with specific genotypes at neighboring SNPs (within a window of 1 Mbp), indicating that particular genetic backgrounds may influence methylation levels differently in ME/CFS patients than in controls. The genomic elements associated with genetic and epigenetic variants characteristic of ME/CFS patients in this study constitute targets for future research. Understanding the molecular mechanisms of genetic-epigenetic interactions of these targets will be key to develop new treatments for ME/CFS, and can serve as a model to understand the molecular basis of related complex diseases.

## Acknowledgements

Funding for this research was provided by operating funds from the Solve ME/CFS Initiative. S.H. was supported by a CIHR Fellowship – Priority Announcement: Myalgic Encephalomyelitis/CFS/FM award number FRN 141047. We would like to thank Dr. Benjamin Hing for helpful discussions.

## Author contributions

P.O.M. designed research; S.H. performed research; S.H. analyzed the data with contributions from W.C.D.V. and D.A.; S.D.V. contributed reagents/materials; S.H. wrote the manuscript with contributions from W.C.D.V., P.O.M., D.A. and S.D.V. All authors read and approved the final manuscript.

## Accession Codes

SNP data will be made available through the NCBI dbSNP database, and methylation data through the NCBI GEO database, upon acceptance for publication.

## References

1. Jason, L. A., Richman, J. A., Rademaker, A. W. & Jordan, K. M. A Community-Based Study of Chronic Fatigue Syndrome. Arch. Intern. Med. 159, 2129–2137 (1999).

2. Jason, L. A., Benton, M. C., Valentine, L., Johnson, A. & Torres-Harding, S. The economic impact of ME/CFS: individual and societal costs. Dyn. Med. 7, 6 (2008).

3. Hornig, M. et al. Distinct plasma immune signatures in ME/CFS are present early in the course of illness. Sci. Adv. 1, e1400121 (2015).

4. Landi, A., Broadhurst, D., Vernon, S. D., Tyrrell, D. L. J. & Houghton, M. Reductions in circulating levels of IL-16, IL-7 and VEGF-A in myalgic encephalomyelitis/chronic fatigue syndrome. Cytokine 78, 27–36 (2016).

5. Brenu, E. W. et al. Role of adaptive and innate immune cells in chronic fatigue syndrome/myalgic encephalomyelitis. Int. Immunol. 26, 233–242 (2014).

6. Brenu, E. W. et al. Immunological abnormalities as potential biomarkers in Chronic Fatigue Syndrome/Myalgic Encephalomyelitis. J. Transl. Med. 9, 81 (2011).

7. Klimas, N. G., Salvato, F. R., Morgan, R. & Fletcher, M. A. Immunologic abnormalities in chronic fatigue syndrome. J. Clin. Microbiol. 28, 1403–1410 (1990).

8. Fluge, Ø. et al. Benefit from b-lymphocyte depletion using the anti-CD20 antibody rituximab in chronic fatigue syndrome. a double-blind and placebo-controlled study. PLoS One 6, (2011).

9. Gaab, J. et al. Low-dose dexamethasone suppression test in chronic fatigue syndrome and health. Psychosom. Med. 64, 311–318 (2002).

10. Sánchez, J. A. & Dorado, D. Intragenomic ITS2 variation in Caribbean seafans. Proceedings of the 11th International Coral Reef Symposium 1383–1387 (2008). at <http://www.nova.edu/ncri/11icrs/proceedings/files/m26-11.pdf>

11. Brothers, L. L. et al. Evidence for extensive methane venting on the southeastern U.S. Atlantic margin. Geology 41, 807–810 (2013).

12. Van Den Eede, F., Moorkens, G., Van Houdenhove, B., Cosyns, P. & Claes, S. J. Hypothalamic-pituitary-adrenal axis function in chronic fatigue syndrome. Neuropsychobiology 55, 112–120 (2007).

13. Visser, J. et al. Increased Sensitivity to Glucocorticoids in Peripheral Blood Mononuclear Cells of Chronic Fatigue Syndrome Patients, Without Evidence for Altered Density or Affinity of Glucocorticoid Receptors. J. Investig. Med. 49, 195–204 (2001).

14. de Vega, W. C., Herrera, S., Vernon, S. D. & McGowan, P. O. Epigenetic modifications and glucocorticoid sensitivity in Myalgic Encephalomyelitis/Chronic Fatigue Syndrome (ME/CFS). BMC Med. Genomics 10, 11 (2017).

15. Schlauch, K. A. et al. Genome-wide association analysis identifies genetic variations in subjects with myalgic encephalomyelitis/chronic fatigue syndrome. Transl. Psychiatry 6, e730 (2016).

16. Rajeevan, M. S., Dimulescu, I., Murray, J., Falkenberg, V. R. & Unger, E. R. Pathway-focused genetic evaluation of immune and inflammation related genes with chronic fatigue syndrome. Hum. Immunol. 76, 553–560 (2015).

17. Smith, A. K., Fang, H., Whistler, T., Unger, E. R. & Rajeevan, M. S. Convergent genomic studies identify association of GRIK2 and NPAS2 with chronic fatigue syndrome. Neuropsychobiology 64, 183–194 (2011).

18. de Vega, W. C., Vernon, S. D. & McGowan, P. O. DNA Methylation Modifications Associated with Chronic Fatigue Syndrome. PLoS One 9, e104757 (2014).

19. Brenu, E. W., Staines, D. R. & Marshall-Gradisnik, S. M. Methylation Profile of CD4+ T Cells in Chronic Fatigue Syndrome/Myalgic Encephalomyelitis. J. Clin. Cell. Immunol. 5, (2014).

20. Viser, J. CD4 T lymphocytes from patients with chronic fatigue syndrome have a decreased interferon gamma production and increased sensitivity to dexamethasone. J. Infect. Dis. 177, 451–454 (1998).

21. Evengard, B., Jacks, A., Pedersen, N. L. & Sullivan, P. F. The epidemiology of chronic fatigue in the Swedish Twin Registry. Psychol. Med. 35, 1317 (2005).

22. Reyes, M. et al. Prevalence and incidence of chronic fatigue syndrome in Wichita, Kansas. Arch. Intern. Med. 163, 1530–1536 (2003).

23. Reeves, W. C. et al. Prevalence of chronic fatigue syndrome in metropolitan, urban, and rural Georgia. Popul. Health Metr. 5, 5 (2007).

24. Fukuda, K. et al. The chronic fatigue syndrome: a comprehensive approach to its definition and study. International Chronic Fatigue Syndrome Study Group. Ann. Intern. Med. 121, 953–959 (1994).

25. Carruthers, B. M. et al. Myalgic Encephalomyelitis / Chronic Fatigue Syndrome: Clinical Working Case Definition, Diagnostic and Treatment Protocols. J. Chronic Fatigue Syndr. 11, 7–36 (2003).

26. Hays, R. D., Sherbourne, C. D. & Mazel, R. M. The rand 36-item health survey 1.0. Health Econ. 2, 217–227 (1993).

27. Karssen, L. C., van Duijn, C. M. & Aulchenko, Y. S. The GenABEL Project for statistical genomics. F1000Research 5, 914 (2016).

28. Purcell, S. et al. PLINK: A tool set for whole-genome association and population-based linkage analyses. Am. J. Hum. Genet. 81, 559–575 (2007).

29. Anderson, C. A. et al. Data quality control in genetic case-control association studies. Nat. Protoc. 5, 1564–1573 (2010).

30. Clarke, G. M. et al. Basic statistical analysis in genetic case-control studies. Nat. Protoc. 6, 121–33 (2011).

31. Patterson, N., Price, A. L. & Reich, D. Population structure and eigenanalysis. PLoS Genet. 2, 2074–2093 (2006).

32. Price, A. L. et al. Principal components analysis corrects for stratification in genome-wide association studies. Nat. Genet. 38, 904–909 (2006).

33. Dunn, O. J. Multiple Comparisons Among Means. J. Am. Stat. Assoc. 56, 52–64 (1961).

34. Holm, S. A Simple Sequentially Rejective Multiple Test Procedure. Scand. J. Stat. 6, 65– 70 (1979).

35. Benjamini, Y. & Hochberg, Y. Controlling the false discovery rate: a practical and powerful approach to multiple testing. J. R. Stat. Soc. B 57, 289–300 (1995).

36. Barrett, J. C., Fry, B., Maller, J. & Daly, M. J. Haploview: Analysis and visualization of LD and haplotype maps. Bioinformatics 21, 263–265 (2005).

37. MacArthur, J. et al. The new NHGRI-EBI Catalog of published genome-wide association studies (GWAS Catalog). Nucleic Acids Res. 45, D896–D901 (2017).

38. Aken, B. L. et al. Ensembl 2017. Nucleic Acids Res. 45, D635–D642 (2017).

39. Arnold, M., Raffler, J., Pfeufer, A., Suhre, K. & Kastenmüller, G. SNiPA: an interactive, genetic variant-centered annotation browser. Bioinformatics 31, 1334–1336 (2015).

40. GTEx Consortium. The Genotype-Tissue Expression (GTEx) project. Nat. Genet. 45, 580–585 (2013).

41. Suhre, K. et al. Connecting genetic risk to disease end points through the human blood plasma proteome. Nat. Commun. 8, 14357 (2017).

42. Hannon, E. et al. Methylation QTLs in the developing brain and their enrichment in schizophrenia risk loci. Nat. Neurosci. 19, 48–54 (2016).

43. Carvalho, B. S., Louis, T. A. & Irizarry, R. A. Quantifying uncertainty in genotype calls. Bioinformatics 26, 242–249 (2010).

44. Scharpf, R. B., Irizarry, R. A., Ritchie, M. E., Carvalho, B. & Ruczinski, I. Using the R Package crlmm for genotyping and copy number estimation. J. Stat. Softw. 40, 1–32 (2011).

45. Machiela, M. J. & Chanock, S. J. LDlink: a web-based application for exploring population-specific haplotype structure and linking correlated alleles of possible functional variants. Bioinformatics 31, 3555–3557 (2015).

46. de Leeuw, C. A., Mooij, J. M., Heskes, T. & Posthuma, D. MAGMA: Generalized Gene-Set Analysis of GWAS Data. PLOS Comput. Biol. 11, e1004219 (2015).

47. Subramanian, A. et al. Gene set enrichment analysis: a knowledge-based approach for interpreting genome-wide expression profiles. Proc. Natl. Acad. Sci. U. S. A. 102, 15545– 15550 (2005).

48. Liberzon, A. et al. The Molecular Signatures Database (MSigDB) hallmark gene set collection. Cell Syst. 1, 417–425 (2015).

49. Aryee, M. J. et al. Minfi: A flexible and comprehensive Bioconductor package for the analysis of Infinium DNA methylation microarrays. Bioinformatics 30, 1363–1369 (2014).

50. Wilhelm-Benartzi, C. S. et al. Review of processing and analysis methods for DNA methylation array data. Br. J. Cancer 109, 1394–402 (2013).

51. Maksimovic, J., Gordon, L. & Oshlack, A. SWAN: Subset-quantile within array normalization for illumina infinium HumanMethylation450 BeadChips. Genome Biol. 13, R44 (2012).

52. Du, P. et al. Comparison of Beta-value and M-value methods for quantifying methylation levels by microarray analysis. BMC Bioinformatics 11, 587 (2010).

53. Johnson, W. E., Li, C. & Rabinovic, A. Adjusting batch effects in microarray expression data using empirical Bayes methods. Biostatistics 8, 118–127 (2007).

54. Jaffe, A. E. et al. Bump hunting to identify differentially methylated regions in epigenetic epidemiology studies. Int. J. Epidemiol. 41, 200–209 (2012).

55. Shabalin, A. A. Matrix eQTL: Ultra fast eQTL analysis via large matrix operations. Bioinformatics 28, 1353–1358 (2012).

56. Kuhn, R. M. et al. The UCSC Genome Browser Database: update 2009. Nucleic Acids Res. 37, D755–D761 (2009).

57. Yu, G., Wang, L.-G., Han, Y. & He, Q.-Y. clusterProfiler: an R package for comparing biological themes among gene clusters. OMICS 16, 284–287 (2012).

58. Ashburner, M. et al. Gene ontology: tool for the unification of biology. The Gene Ontology Consortium. Nat. Genet. 25, 25–29 (2000).

59. Gene Ontology Consortium. Gene Ontology Consortium: going forward. Nucleic Acids Res. 43, D1049–D1056 (2015).

60. Kanehisa, M. & Goto, S. KEGG: kyoto encyclopedia of genes and genomes. Nucleic Acids Res. 28, 27–30 (2000).

61. Kanehisa, M., Goto, S., Sato, Y., Furumichi, M. & Tanabe, M. KEGG for integration and interpretation of large-scale molecular data sets. Nucleic Acids Res. 40, D109–14 (2012).

62. Yu, G. & He, Q.-Y. ReactomePA: an R/Bioconductor package for reactome pathway analysis and visualization. Mol. Biosyst. 12, 477–479 (2016).

63. Yu, G., Wang, L.-G., Yan, G.-R. & He, Q.-Y. DOSE: an R/Bioconductor package for disease ontology semantic and enrichment analysis. Bioinformatics 31, 608–609 (2015).

64. Curriu, M. et al. Screening NK-, B- and T-cell phenotype and function in patients suffering from Chronic Fatigue Syndrome. J. Transl. Med. 11, 68 (2013).

65. Hardcastle, S. L. et al. Characterisation of cell functions and receptors in Chronic Fatigue Syndrome/Myalgic Encephalomyelitis (CFS/ME). BMC Imunol. 16, 35 (2015).

66. Little, T. J., Horowitz, M. & Feinle-Bisset, C. Role of cholecystokinin in appetite control and body weight regulation. Obes. Rev. 6, 297–306 (2005).

67. Lignon, M. F., Bernad, N. & Martinez, J. Cholecystokinin receptors in cells of the immune system. Ann. N. Y. Acad. Sci. 713, 334–337 (1994).

68. Weiland, T. J., Voudouris, N. J. & Kent, S. CCK(2) receptor nullification attenuates lipopolysaccharide-induced sickness behavior. Am. J. Physiol. Regul. Integr. Comp. Physiol. 292, R112–R123 (2007).

69. Morris, G., Anderson, G., Galecki, P., Berk, M. & Maes, M. A narrative review on the similarities and dissimilarities between myalgic encephalomyelitis/chronic fatigue syndrome (ME/CFS) and sickness behavior. BMC Med. 11, 64 (2013).

70. Chung, S. et al. Identification of preoptic sleep neurons using retrograde labelling and gene profiling. Nature 545, 477–481 (2017).

71. Zhang, J.-G. et al. Cholecystokinin octapeptide regulates the differentiation and effector cytokine production of CD4(+) T cells in vitro. Int. Immunopharmacol. 20, 307–315 (2014).

72. Geibel, M. et al. Ablation of TrkB signalling in CCK neurons results in hypercortisolism and obesity. Nat. Commun. 5, 3427 (2014).

73. Spada, J. et al. Genome-wide association analysis of actigraphic sleep phenotypes in the LIFE Adult Study. J. Sleep Res. 25, 690–701 (2016).

74. Guo, C. et al. Myeloid-derived suppressor cells have a proinflammatory role in the pathogenesis of autoimmune arthritis. Ann. Rheum. Dis. 75, 278–285 (2016).

75. Graves, M. C. et al. Methylation differences at the HLA-DRB1 locus in CD4+ T-Cells are associated with multiple sclerosis. Mult. Scler. 20, 1033–41 (2014).

76. Maes, M. Inflammatory and oxidative and nitrosative stress pathways underpinning chronic fatigue, somatization and psychosomatic symptoms. Current opinion in psychiatry 22, (2009).

77. Morris, G. et al. Nitrosative Stress, Hypernitrosylation, and Autoimmune Responses to Nitrosylated Proteins: New Pathways in Neuroprogressive Disorders Including Depression and Chronic Fatigue Syndrome. Mol. Neurobiol. 54, 4271–4291 (2017).

78. Naviaux, R. K. et al. Metabolic features of chronic fatigue syndrome. Proc. Natl. Acad. Sci. 201607571 (2016). doi:10.1073/pnas.1607571113

79. Heyn, H. et al. DNA methylation contributes to natural human variation. Genome Res. 23, 1363–1372 (2013).

80. Glossop, J. R. et al. Genome-wide DNA methylation profiling in rheumatoid arthritis identifies disease-associated methylation changes that are distinct to individual T- and B-lymphocyte populations. Epigenetics 9, 1228–1237 (2014).

81. Li, J.-P. et al. The phosphatase JKAP/DUSP22 inhibits T-cell receptor signalling and autoimmunity by inactivating Lck. Nat. Commun. 5, 1–13 (2014).

